# Effects of a high cholesterol diet on *Drosophila* chill tolerance are highly context-dependent

**DOI:** 10.1101/2023.09.09.556984

**Authors:** Mitchell C. Allen, Marshall W. Ritchie, Mahmoud I. El-Saadi, Heath A. MacMillan

## Abstract

Chill susceptible insects are thought to be injured through different mechanisms depending on the duration and severity of chilling. While chronic chilling causes “indirect” injury through disruption of metabolic and ion homeostasis, acute chilling is suspected to cause “direct” injury, in part through phase transitions of cell membrane lipids. Dietary supplementation of cholesterol can reduce acute chilling injury in *Drosophila melanogaster*, but the generality of this effect and the mechanisms underlying it remain unclear. To better understand how and why cholesterol has this effect, we assessed how a high cholesterol diet and thermal acclimation independently and interactively impact several measures of chill tolerance in both male and female flies. Cholesterol supplementation positively affected tolerance to acute chilling in warm-acclimated flies (as reported previously). Conversely, feeding on the high-cholesterol diet negatively affected tolerance to chronic chilling in both cold and warm acclimated flies, as well as tolerance to acute chilling in cold acclimated flies. Cholesterol had no effect on the ability of flies to remain active in the cold or recover movement after a cold stress. Our findings support the idea that dietary cholesterol reduces mechanical injury to membranes caused by direct chilling injury, and that acute and chronic chilling are associated with distinct mechanisms of injury. Feeding on a high-cholesterol diet may interfere with mechanisms involved in cold acclimation, leaving cholesterol augmented flies more susceptible to chilling injury under some conditions.

**Highlights:**

- Cholesterol improves cold shock tolerance of warm-acclimated flies
- Cold acclimation and chronic cold instead lead to negative effects of cholesterol on chill tolerance
- Cholesterol did not affect the ability of flies to remain active in the cold
- Both sexes were similarly affected by a high cholesterol diet

## Introduction

Many ectothermic insects living in cold climates can either avoid or tolerate extracellular ice formation, and thereby prevent injury or death at low temperatures (Lee, 1991). Despite the effectiveness of these strategies in mitigating injury related to freezing, many insects are ‘chill susceptible’, and are injured at low temperatures through mechanisms unrelated to freezing (Bale, 1996; Overgaard and Macmillan, 2017).

Tolerance to chilling in insects can be quantified in multiple ways. For example, chill coma onset (CCO) is the temperature at which an insect loses all capacity for movement and is indicative of the thermal limits of neuromuscular function (MacMillan and Sinclair, 2011; Robertson et al., 2023). It can be measured by gradually reducing ambient temperature and recording the temperature at which all movement ceases. Additionally, cold tolerance can be evaluated by performing injury/survival experiments, where individuals are exposed to a low temperature (typically below CCO) for a pre-determined period before they are rewarmed and the number or proportion of individuals who are injured or killed is counted (Rozsypal et al., 2021; Shreve et al., 2007; Sinclair and Chown, 2003). The degree of injury inflicted during the exposure can also be determined by grading the physical condition of insects on a multi-point scale, rather than assigning them a binary (alive/dead) survival grade (El-Saadi et al., 2020; MacMillan et al., 2014). Such injury assays performed following either acute or chronic chilling can provide useful insights into the proximate and ultimate mechanisms of chilling injury. If the cold stress is mild or brief enough to not cause lethal injury, chill coma recovery time (CCRT) is a measure of time required for an insect to stand after being removed from the cold stress to a permissible temperature (Gibert et al., 2001).

Many species of insects can alter their thermal tolerance within their lifetime; this phenotypic plasticity allows insects to adjust to temperature changes over multiple timescales, such as changing seasons or diurnal temperature changes (Shearer et al., 2016). In the laboratory, pre-exposure to a low temperature that does not cause injury can improve tolerance to a future cold stress (Czajka and Lee, 1990; Rako and Hoffmann, 2006). An insect can respond rapidly to a temperature change through rapid cold-hardening (RCH), a response that occurs within minutes or a few hours of cold exposure (Colinet and Hoffmann, 2012; Czajka and Lee, 1990; Sinclair and Roberts, 2005; Teets et al., 2019). If chilling occurs over a longer period (e.g. days or weeks), the response is typically termed cold acclimation (Gerken et al., 2015; Koštál et al., 2011; Rako and Hoffmann, 2006).

The mechanisms underlying different chill tolerance traits are related, but distinct, meaning these traits can vary independently (Davis et al., 2021). While CCO is caused by the silencing of neuromuscular signaling from temperature effects on ionoregulation in the central nervous system (Andersen and Overgaard, 2019), CCRT relates to recovery of this local ion balance as well as restoration of systemic ion homeostasis via the renal system (MacMillan et al., 2012). The causes of chilling injury are thought to vary depending on the nature (intensity and duration) of the cold exposure (Sinclair and Roberts, 2005). Extended periods of chilling (chronicchilling) leads to progressive disruption of ion/water balance resulting in hemolymph hyperkalemia, that drives cell depolarization and cell death (Bayley et al., 2018; Carrington et al., 2020; MacMillan, 2019; Macmillan et al., 2015). By contrast, severe acute chilling (a shorter exposure to a more severe low temperature) is thought to instead cause cell death through temperature-induced permeabilization of cell lipid bilayers via a phase transition of membrane lipids (Turnock and Bodnaryk, 1993, 1991).

Cold acclimation can alter chill coma onset temperature through effects on ionoregulatory processes in the nervous system (Des Marteaux et al., 2018; Himmel et al., 2021; MacMillan et al., 2015) and can improve chill tolerance in part by mitigating hyperkalemia in the cold via plasticity of the renal system (Andersen and Overgaard, 2020; Yerushalmi et al., 2018) and the effects of hyperkalemia on cell survival (Andersen et al., 2017). While RCH can also improve cold tolerance, it is driven at least in part by mechanisms unrelated to those associated with cold acclimation (Teets and Denlinger, 2013). Unlike acclimation, RCH is thought to occur mostly at the cellular level, having been demonstrated in isolated cells *in vitro* (Nadeau and Teets, 2020; Yi and Lee, 2004).

Thermal plasticity (like acclimation or RCH) enables animals to alter the composition of their cell membranes by inserting or removing various lipids, proteins, or sterols which assists them in adjusting to prevailing environmental conditions. This phenomenon, called homeoviscous adaptation, has been observed in a wide range of ectotherms (Koštál et al., 2003; Logue et al., 2000; Snyder and Hennessey, 2003). Cholesterol is involved in homeoviscous adaptation and plays an important role in regulating the structure, function, and fluidity of cell membranes and membrane bound proteins (Cooper, 1978; Klaiss-Luna and Manrique-Moreno, 2022; Yeagle, 1991). Cholesterol also provides a structural base for the formation of lipid rafts, which provide concentrated regions of glycosphingolipids and protein receptors on the surface of cell membranes (Lee et al., 2021; Thomas et al., 2004). These microdomains play an important role in membrane fluidity as well as cell signalling and membrane trafficking (Komatsuya et al., 2020; Simons and Ikonen, 1997). While the role of cholesterol in modulating membrane fluidity and stability is well documented, our understanding of its role in insect chill tolerance is incomplete.

Dietary augmentation of cholesterol has been shown to improve survival following acute cold exposure in *D. melanogaster* (Shreve et al., 2007), but what other measures of cold tolerance are affected by cholesterol, and how, remains unclear. Analysis of whole-fly membrane preparations suggests that cholesterol acquired from the diet is integrated into cell membranes (Shreve et al., 2007). Since insects cannot produce cholesterol endogenously, any cholesterol incorporated into their cell membranes must be acquired through diet (Igarashi et al., 2018). A high-cholesterol diet and RCH pre-treatment also have additive positive effects on cold tolerance, suggesting that additional cholesterol inserted into cell membranes interacts with mechanisms underlying RCH (Shreve et al., 2007). Long-term acclimation, which was not investigated in the same study, has also be associated with homeoviscous adaptation in *Drosophila* (Overgaard et al., 2008), but how cholesterol impacts cold acclimation is unclear.

Investigating the interactive effects of cholesterol augmentation and cold acclimation on cold tolerance could assist in determining the specific role cholesterol plays in thermal plasticity. The current evidence suggesting that cholesterol supplementation alters insect cold tolerance is limited to acute cold exposure (Shreve et al., 2007), which suggest that it might alter chill injury by mitigating mechanical injury to cell membranes. Given that injury caused by acute and chronic cold exposures are thought to be mechanistically distinct (through direct and indirect mechanisms), it is important to consider how the effects of high cholesterol feeding might differ in flies exposed to cold treatments of varying intensity. Investigating the effects of cholesterol on tolerance to chronic chilling, for example, would assist in determining its impact on ion and water balance in the cold, which is heavily dependent on the membrane environment in which ion-motive transporters sit. By contrast, identifying the effects of cholesterol on chill coma phenotypes may reveal its impacts on the maintenance of nerve and muscle excitability in the cold. At the same time, insight into how cholesterol may affect chill tolerance differently between sexes is also lacking. Sex is an important factor to consider because cholesterol is also important to insect reproduction; female insects require high levels of cholesterol for embryogenesis, and males incorporate cholesterol as a structural component of sperm cells (Behmer and Grebenok, 1998; Jing and Behmer, 2020).

In this study, we test the hypothesis that cholesterol and cold acclimation additively or synergistically improve all aspects of insect chill tolerance. We reared male and female flies on either a standard cornmeal-based (control) diet or a cholesterol augmented diet (Table 1) and acclimated them to either 25°C or 15°C. Chill tolerance traits (CCO, CCRT, and survival following acute and chronic chilling) were assayed using both warm-acclimated and cold-acclimated flies. Fruit flies can take three generations to adapt to a change in diet, over which time their rates of reproduction are reduced (Mayntz and Toft, 2001). The impacts of additional cholesterol on chill tolerance could therefore vary within the first few generations of feeding. For this reason, we also measured CCO and acute chilling tolerance over the course of four generations using warm-acclimated flies. All other traits were assayed after at least four generations of a high-cholesterol diet (Shreve et al., 2007).

**Table 1:** Recipe of Bloomington diet (control diet) with quantities per 1 L of food.

| Ingredient | Quantity |
| --- | --- |
| Distilled water | 0.92 L |
| Yellow cornmeal | 67.1 g |
| Inactive yeast | 15.9 g |
| *Drosophila* agar | 8 g |
| Corn syrup | 70 g |
| Light dry malt | 42.3 g |
| Soy flour | 9.2 g |
| Propionic acid | 4.4 mL |

## Materials and Methods

### Animal husbandry

The line of *Drosophila melanogaster* in this experiment originated from 35 isofemale lines that were gathered from London and Niagara, Ontario, Canada (Marshall and Sinclair, 2010). The flies were reared in 200 mL plastic bottles containing 50 mL of a Bloomington diet (control diet; Table 1). The cholesterol-augmented diet had 1 mg of cholesterol added per mL of the standard diet (as in Shreve et al., 2007).

Flies were kept in an incubator (model MIR-154-PA; Panasonic Holdings, Kadoma, Osaka, Japan) at 25°C in a 12 h:12 h light/dark cycle. New eggs were gathered by transferring ∼100 adult flies into a new bottle containing fresh food and were left for 2-3 h to lay eggs before being removed (roughly 80-120 eggs per bottle). Once adult flies had emerged, they were transferred to bottles with fresh food. Flies were kept on the cholesterol-augmented or control diet for the entire duration of rearing. Three days after emerging, experimental flies were sorted by sex under a microscope while lightly anesthetized by CO_2_ for no more than 10 min (MacMillan et al., 2017; Nilson et al., 2006). Sorted flies were transferred in groups of 10 to 50 mL vials containing 10 mL of their assigned diet.

### Acute chilling tolerance and cholesterol feeding over four generations

25°C acclimated adult male and female flies reared on the control and cholesterol-augmented diets were used to assess how cholesterol influences acute chilling tolerance (i.e. that related to “direct” chilling injury). The cholesterol augmented flies used for the initial experimental group were the first generation to be reared on the cholesterol diet. In total, 30 males and 30 females from each diet were individually placed in 3.7 mL screw-top vials in sealed plastic bags and submerged in a refrigerated circulating bath (model AP28R-30; VWR international, Radnor, PA, USA) containing a 1:1 v/v mixture of ethylene glycol and water. Flies were exposed to -4°C or -5°C for 1 h, then returned to 1.5 mL microcentrifuge tubes containing ∼0.5 mL of their respective diet to recover at 25°C (O’Neill et al., 2021). The physical condition of the flies was assessed after 4 h and 24 h of recovery using a 4-point scale based on their behaviour, as adapted from MacMillan et al., (2014). The scoring system was: 4, able to walk and fly; 3, able to stand but unable to fly; 2, unable to stand but exhibit some movement; 1, no movement. Flies from the same lineage which were not sorted by sex were used to maintain populations of flies continually raised on both diets and continue the experiment over four subsequent generations.

### Acute and chronic chilling tolerance in combination with thermal acclimation

For experiments involving cold acclimation, flies that had been fed on cholesterol for at least four generations (matching the methods of Shreve et al., 2007) were raised as described above. Flies were sorted under CO_2_ anaesthesia 3 d post adult emergence as above. At this time, roughly half of the flies were pseudo-randomly selected to be left at 25°C, and the other half was transferred to 15°C in a 12 h:12 h light/dark cycle to acclimate for four days in groups of 10 flies within 50 mL vials containing 10 mL of the assigned diet. Both warm-(25°C) and cold-acclimated (15°C) flies were all 7 d post adult ecdysis at the time of experiments.

To test how a high cholesterol diet interacted with cold acclimation to influence chilling injury, we exposed both warm- and cold-acclimated flies to a more severe cold stress that would induce quantifiable injury in the cold-acclimated flies and allow us to confirm expected improvements in chilling tolerance typically associated with cold acclimation. Based on preliminary trials we decided on the following: For the acute chilling assay, 15 replicate male and female adult flies from both diets and acclimations were placed in 3.7 mL screw top-vials in sealed plastic bags and submerged in glycol at -5°C for 3 hr. For the chronic chilling assay, 15 replicate male and female adult flies from both diets and acclimations were placed in 5 mL screw top-vials in sealed plastic bags and submerged in a mixture of ice and water for 24 h. Immediately after the treatment, the flies were transferred individually into 1.5 mL microcentrifuge tubes containing ∼0.5 mL of their respective diet. As before, the physical condition of the flies was assessed after 4 h and 24 h of recovery at 25°C using the 4-point scale described for the acute chilling experiment.

### Chill coma onset

Chill coma onset temperature (CCO) was measured in flies raised at 25°C and that remained at 25°C as adults. Both males and females were raised on either a control or the cholesterol-augmented diet, and this process was repeated over four generations of flies feeding on the control and cholesterol-augmented diets as for the acute cold tolerance experiment above. For each generation, twenty replicate flies from each treatment were transferred individually to 3.7 mL glass screw-top vials (MacMillan et al., 2017) The vials containing the flies were placed in a custom rack that was submerged in a bath containing a mixture of ethylene glycol and water (1:1 v/v) (MacMillan et al., 2017). The temperature was held at 20°C for 10 min and then linearly decreased by 0.1°C/min. Observations of the flies started when the temperature was below 10°C, and once flies had stopped moving, they were stimulated by tapping on the vial with a hard plastic rod. The CCO was recorded as the temperature at which a fly ceased to move with no response to the tapping (Gibert and Huey, 2001). This process was repeated over four generations of flies feeding on the control and cholesterol-augmented diets.

### Chill coma onset and recovery following thermal acclimation

To quantify CCO following both cholesterol feeding and thermal acclimation, twenty replicate flies from each treatment were transferred individually to 3.7 mL glass screw-top vials, and CCO was recorded as described for the previous CCO assay across multiple generations. Chill coma recovery time (CCRT) was measured by placing 20 replicate flies from control and cholesterol-augmented diet treatments in 3.7 mL screw-top vials in a sealed plastic bag that was submerged in a mixture of ice and water (0°C) for 6 h. The flies were then removed from the bath and arranged on a counter at room temperature (22°C) for observation. The time it took each fly to stand upright was recorded as CCRT (David et al., 1998; Lachenicht et al., 2010).

### Statistical analyses

All analyses were conducted using R software (version 4.2.2; R core team, 2022). The independent and interactive effects of feeding on a high cholesterol diet and cold acclimation on cold tolerance was determined using generalized linear models (GLM) for CCO and CCRT. Survival data was analyzed using ordinal logistic regressions (MASS, RStata packages). For each analysis, we started with a saturated model and removed non-significant interaction using Akaike’s information criterion (AIC) to identify the best fitting model.

## Results

In most cases both male and female warm acclimated flies feeding on the high-cholesterol diet were better able to tolerate acute chilling compared to the control flies, although the magnitude of the effect of diet was dependant on sex and recovery time (a three-way interaction; *P_Sex*Diet*Recovery_* <0.001; Fig. 2; Table 2). Generation interacted with other factors to influence injury scores (Table 2), but these effects explained a relatively small portion of the variance in the dataset and there was no clear trend of scores improving or worsening over subsequent generations in the cholesterol augmented flies. While significantly more variance is attributed to diet in the fourth generation than the aggregate of all of the generations, we note that this reflects idiosyncratic changes among generations, rather than a continuous trend in the effects of cholesterol (Table 2; Fig. 1). As a result, we decided to aggregate the data across generations in Figure 2. We note that similar trends were observed when performing statistical analyses using only the 4^th^ generation of flies (Table S1). Female flies suffered from further injury after several hours of recovery (*P_Sex*Recovery time_* < 0.001; Table 2), so differences in scores associated with diet were better resolved after 4 h of recovery in females, and 24 h in males, who were also generally more resistant to the cold exposure (*P_Sex_*<0.001; Table 2).

**Figure 1.**
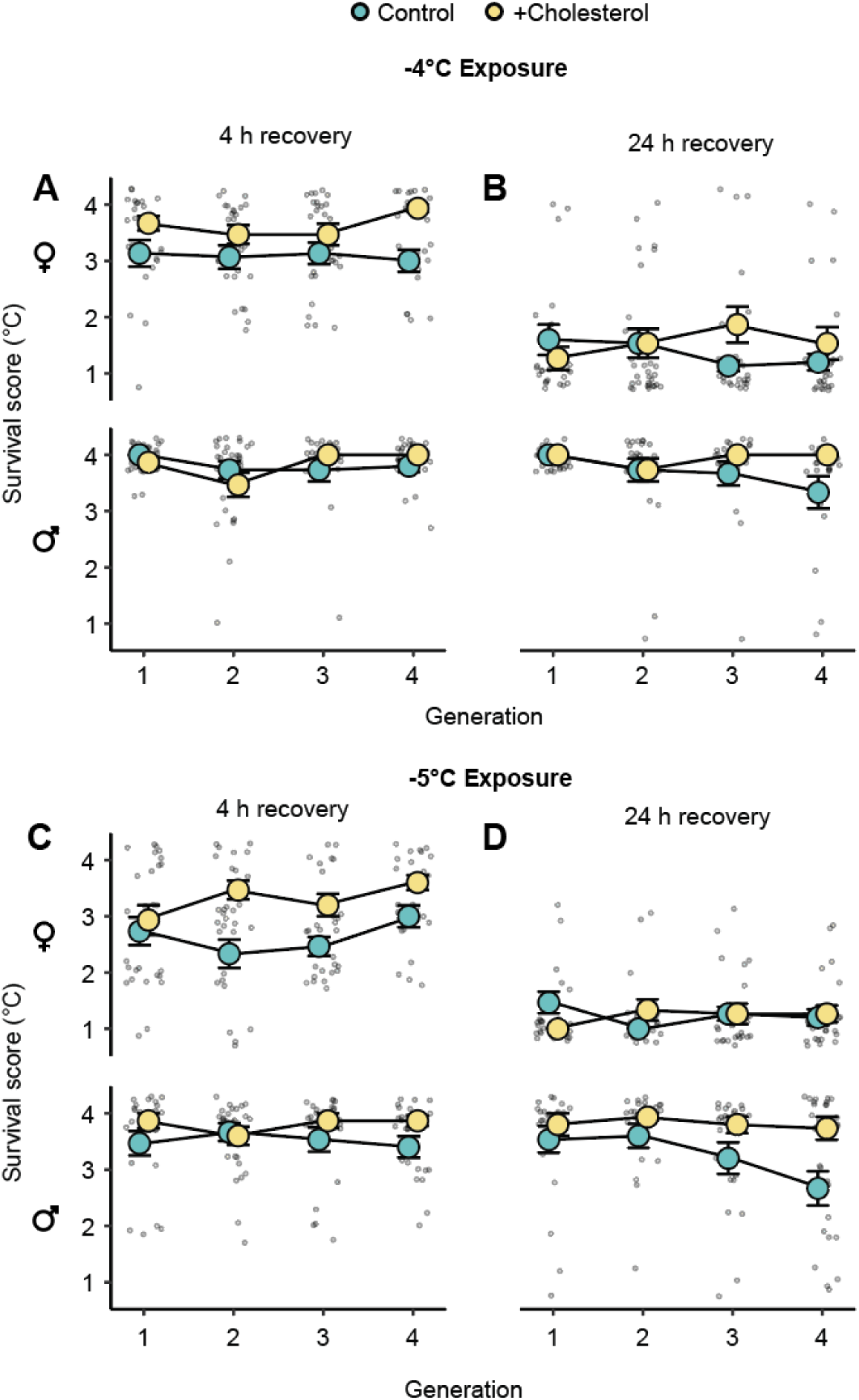
Cholesterol supplementation typically improves tolerance to acute chilling in both male and female *Drosophila melanogaster* across four generations of feeding. Flies were exposed directly to -4°C (A, B) or -5°C (C, D) before being removed to 25°C for recovery. The same flies were scored after recovery for 4 h (A, C) and 24 h (B, D) at 25°C. Output from ordinal logistic regressions. for this experiment are presented in Table 2. Large points with error bars represent the mean ± sem, and small points represent scores of individual flies. Error bars that are not visible are obscured by the symbols.

**Figure 2:**
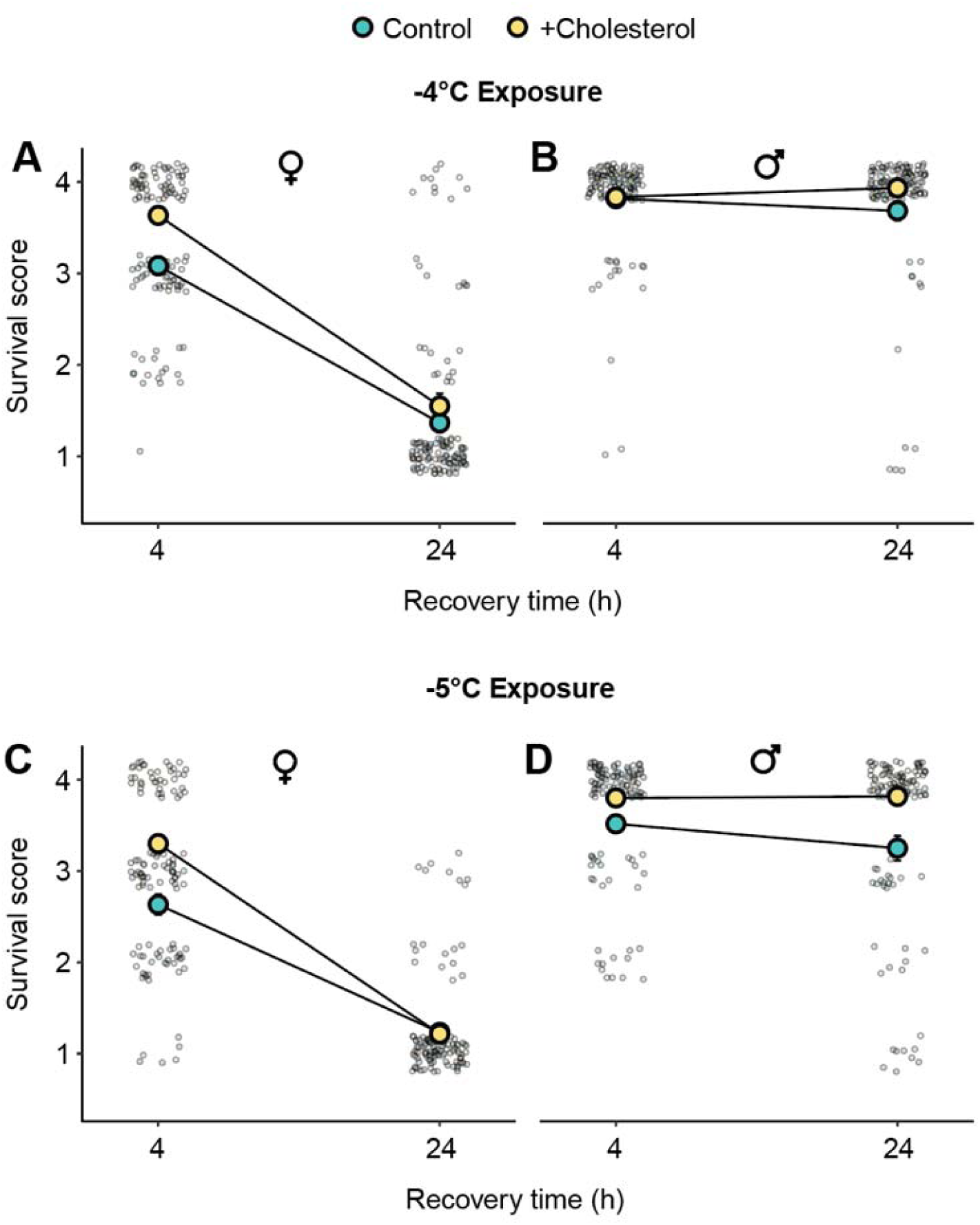
Aggregate data across four generations of flies supports that cholesterol typically protects against direct chilling injury in warm-acclimated flies. Injury scores assigned to male and female flies fed either control or high-cholesterol diets and exposed to acute chilling to -4°C (A,B) or -5°C (C,D) for 1 h. Output of ordinal logistic regression for this dataset are presented in Table 2. Large points with error bars represent the mean ± sem, and small points represent scores of individual flies. Data was aggregated over four generations. Error bars that are not visible are obscured by the symbols.

**Table 2:** Output from ordinal logistic regression testing the effects of cholesterol supplementation (diet), sex, exposure temperature, and recovery time (and retained interactions) on injury scores following acute cold stress (-4°C or -5°C) in *Drosophila melanogaster*. The best fitting model for this dataset (based on AIC) includes a 3-way interaction between sex, diet, and recovery time. Odds ratios (OR) and 95% confidence intervals (95% CI) were calculated for factors in brackets. Main effects and interactions in boldface were statistically significant (P < 0.05).

| Factor | OR | 95% CI | P |
| --- | --- | --- | --- |
| **Sex (male)** | **15.21** | **6.07, 39.25** | **< 0.001** |
| Diet (cholesterol) | 0.99 | 0.43, 2.26 | 0.984 |
| **Recovery time (24 h)** | **0.08** | **0.03, 0.20** | **< 0.001** |
| Generation | 0.97 | 0.77, 1.23 | 0.820 |
| **Exposure temperature (-4°C)** | **2.36** | **1.75, 3.22** | **< 0.001** |
| Sex (male) : Diet (cholesterol) | 0.52 | 0.23, 1.21 | 0.125 |
| **Sex (male) : Recovery time (24 h)** | **19.62** | **8.48, 46.12** | **< 0.001** |
| Sex (male) : Generation | 0.77 | 0.58, 1.03 | 0.073 |
| **Diet (cholesterol) : Recovery time (24 h)** | **0.30** | **0.13, 0.67** | **0.003** |
| **Diet (cholesterol) : Generation** | **1.69** | **1.23, 2.23** | **< 0.001** |
| **Recovery time (24 h) : Generation** | **0.71** | **0.54, 0.94** | **0.018** |
| **Sex (male) : Diet (cholesterol) : Recovery time (24 h)** | **10.89** | **2.90, 43.00** | **< 0.001** |

As with acute cold tolerance, the costs and benefits of feeding on the high cholesterol diet on chill coma onset temperature (CCO) varied among generations relative to the control diet (*P_Diet*Generation_* = 0.014, Table S2), but overall, CCO did not improve with cholesterol feeding across four generations (Fig. S1).

In clear contrast to what was observed in warm acclimated flies, cold-acclimated flies scored worse following acute cold stress when they were fed the high cholesterol diet compared to the control diet, although this was not reflected in the statistical analyses, owing perhaps to the large spread of individual scores for this dataset (Fig. 3A, B; Table 3). Overall however, the cholesterol diet was found independently to significantly lower the survival score (*P_Diet_* < 0.001). Both males and females experienced a similar negative effect from the cholesterol diet, specifically when they were cold acclimated (*P_Sex*Acclimation_*= 0.586).

**Figure 3:**
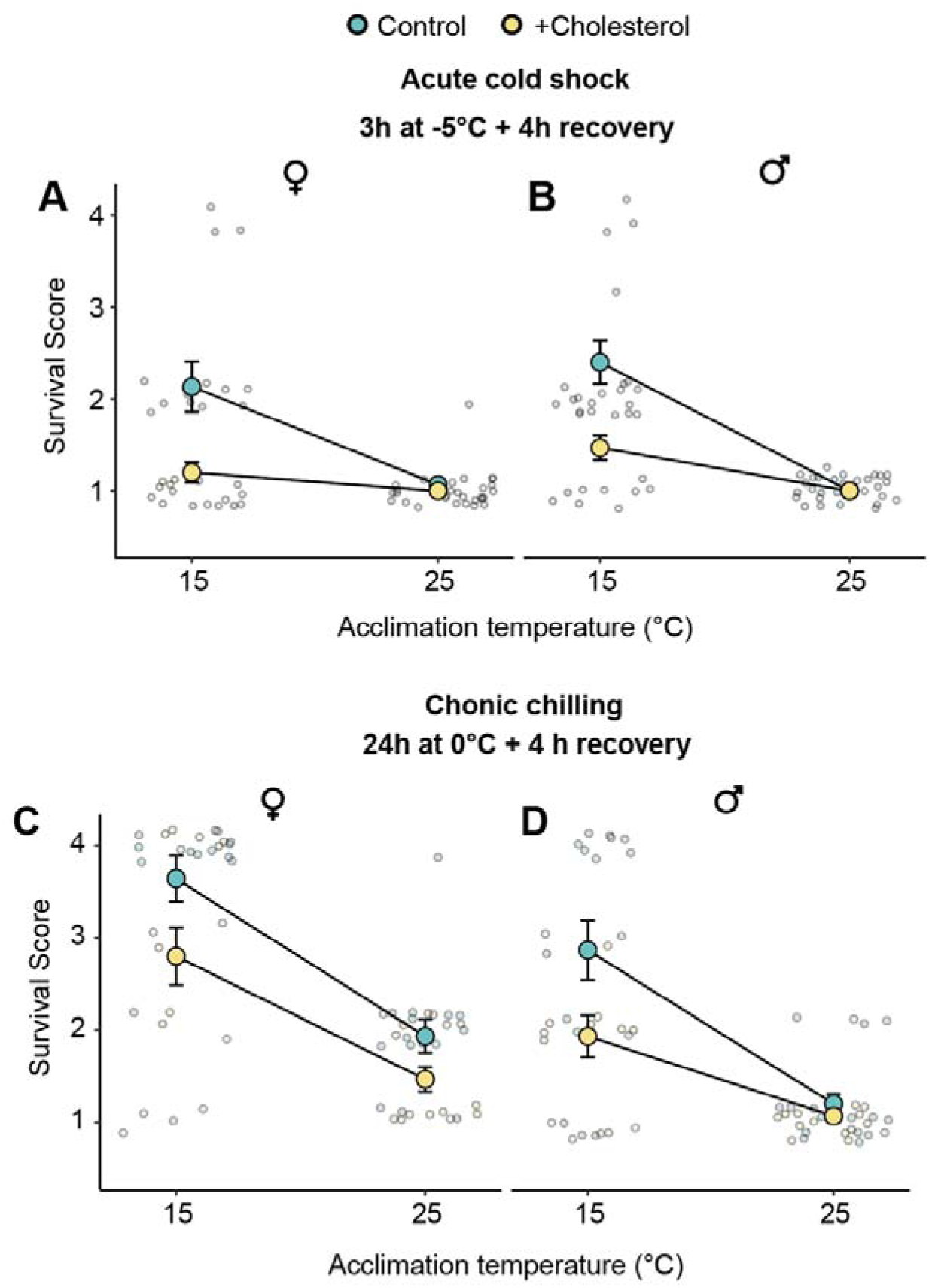
Cold acclimation reverses the protective effects of cholesterol. Injury scores assigned to warm- and cold-acclimated male and female *Drosophila melanogaster* following exposure to -5°C for 3 h. Outputs from ordinal logistic regressions for these plots are presented in Table 3. Large points with error bars represent the mean ± sem, and small points represent scores of individual flies. Error bars that are not visible are obscured by the symbols.

**Table 3:**
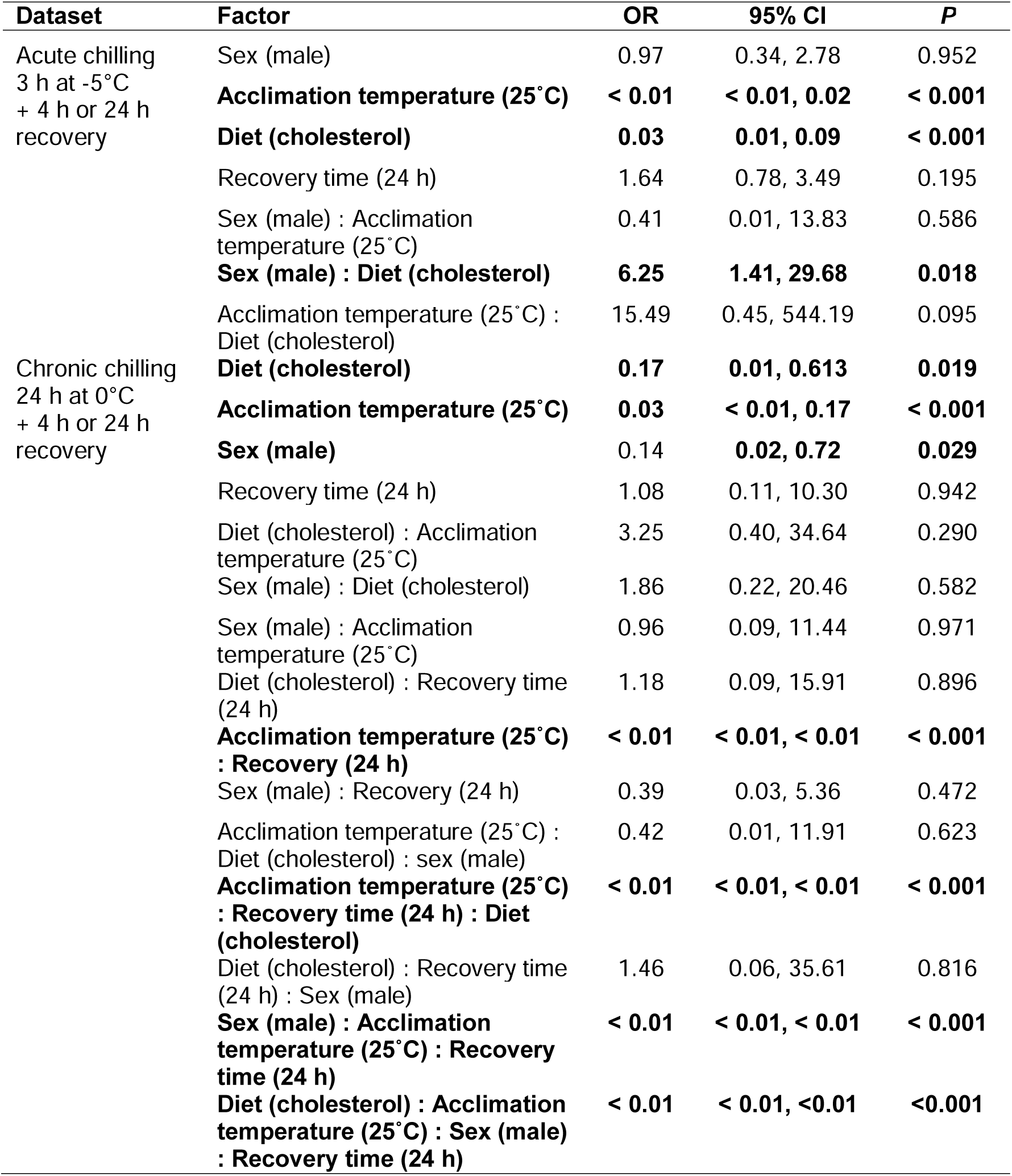
Output from ordinal logistic regressions testing the effects of acclimation temperature, cholesterol supplementation (diet), sex, and recovery time (and retained interactions) on chilling injury scores following either acute or chronic chilling. Best fitting models for these datasets (based on AIC) included 3-way interactions. Odds ratios and 95% confidence intervals were calculated for factors in brackets. Main effects and interactions in boldface were statistically significant (*P* < 0.05).

Flies who were fed the high cholesterol diet were also more injured than flies fed the control diet following a chronic cold exposure, regardless of acclimation status (*P_Diet_* < 0.001; Fig. 3 C, D; Table 3). Again, the negative effect of feeding on the cholesterol diet was similar between sexes in the cold acclimated group (*P_Sex*Acclimation_* = 0.586, Table 3), however males were more injured overall than females by the chronic cold exposure (*P_sex_*= 0.029). Regardless of the diet, warm acclimated flies were severely injured during the cold stress in this experiment, which was expected. These treatments were chosen specifically to resolve the effect of cholesterol on cold-acclimated flies and confirm that cold acclimation improved cold tolerance.

To examine how cholesterol and acclimation influenced other common metrics of chill tolerance, we quantified chill coma onset temperature (CCO) and recovery time (CCRT) in warm and cold-acclimated flies fed control and the cholesterol-augmented diet after more than four generations. Cold acclimation strongly reduced CCO (P_Acclimation_ < 0.001; Fig. 4; Table 4), but was not improved by the high cholesterol diet (*P_Diet_* = 0.683; Fig. 4A, B; Table 4). Diet and acclimation status also did not interact to influence CCO (*P_Diet*Acclimation_*= 0.880, Fig. 4A, B; Table 4), and female flies had higher CCO than males (*P*_sex_ = 0.031; Fig. 4A, B). Similarly, CCRT was unaffected by the cholesterol diet (*P_Diet_* = 0.961 Fig. 4C, D), but cold acclimation significantly reduced CCRT (*P_Acclimation_* < 0.001; Fig. 4C, D), and this benefit did not interact with the high cholesterol diet (Fig. 4 C, D; *P_Diet*Acclimation_* = 0.842).

**Figure 4:**
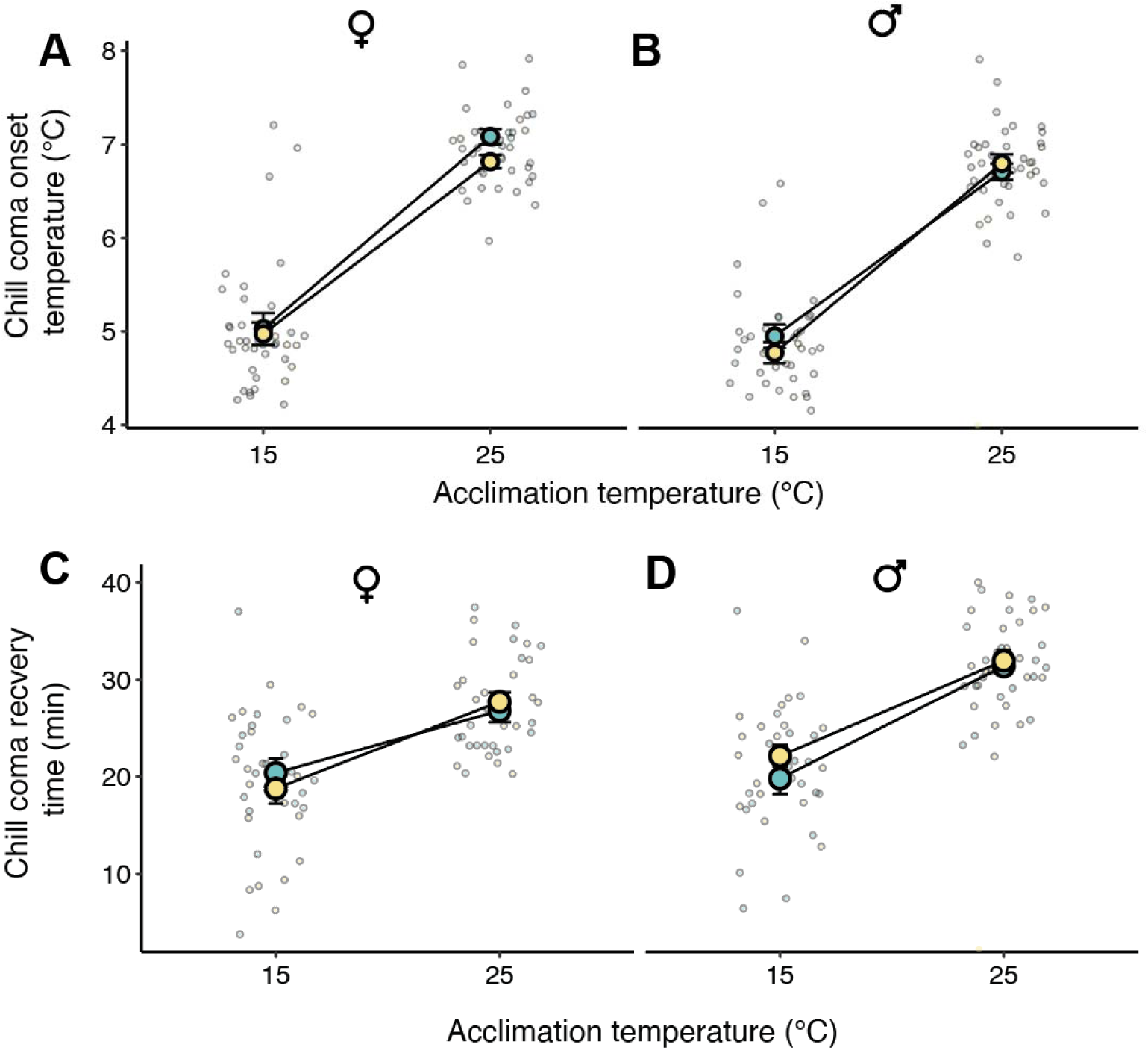
A cholesterol-augmented diet has no effect on chill coma phenotypes regardless of acclimation status. Chill coma onset temperatures (A, B) and chill coma recovery times (C, D) of female and male *D. melanogaster* acclimated to 25°C or 15°C and fed on a control or cholesterol-augmented diet. Large points with error bars represent the mean ± sem, and small points represent scores of individual flies. Outputs from generalized linear models are presented in Table 4. Error bars that are not visible are obscured by the symbols.

**Table 4:**
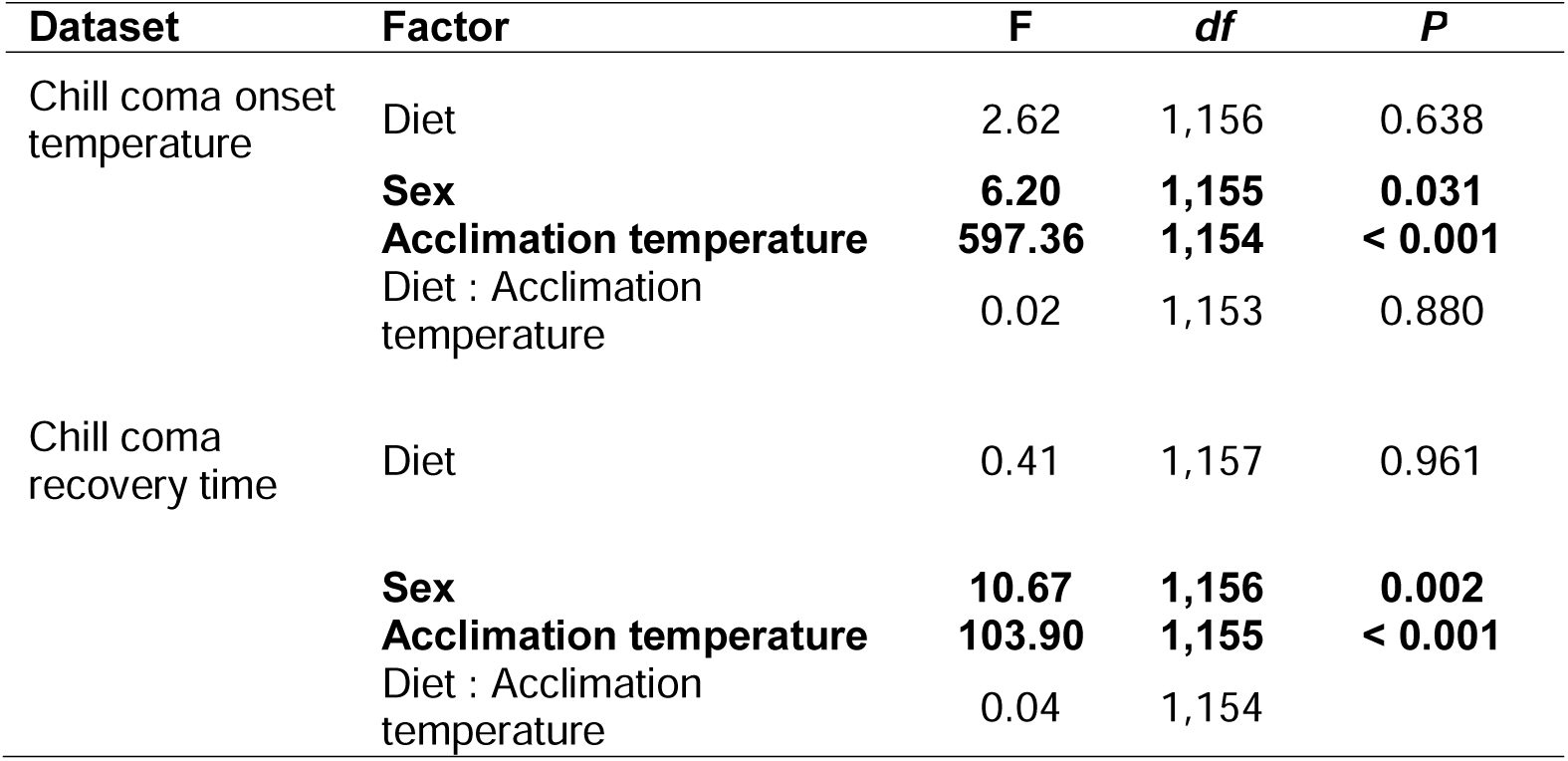
Output from generalized linear model testing the effects of acclimation temperature, cholesterol supplementation (diet), and sex (and retained interactions) on chill coma onset temperature (CCO) and chill coma recovery time (CCRT) in *Drosophila melanogaster*. The best fitting models for these datasets (based on AIC) include non-significant 2-way interactions between diet and acclimation temperature, and the majority of variance in both cases is explained by main effects of thermal acclimation on the chilling tolerance traits. Main effects and interactions in boldface were statistically significant (P < 0.05).

**Figure 5.**
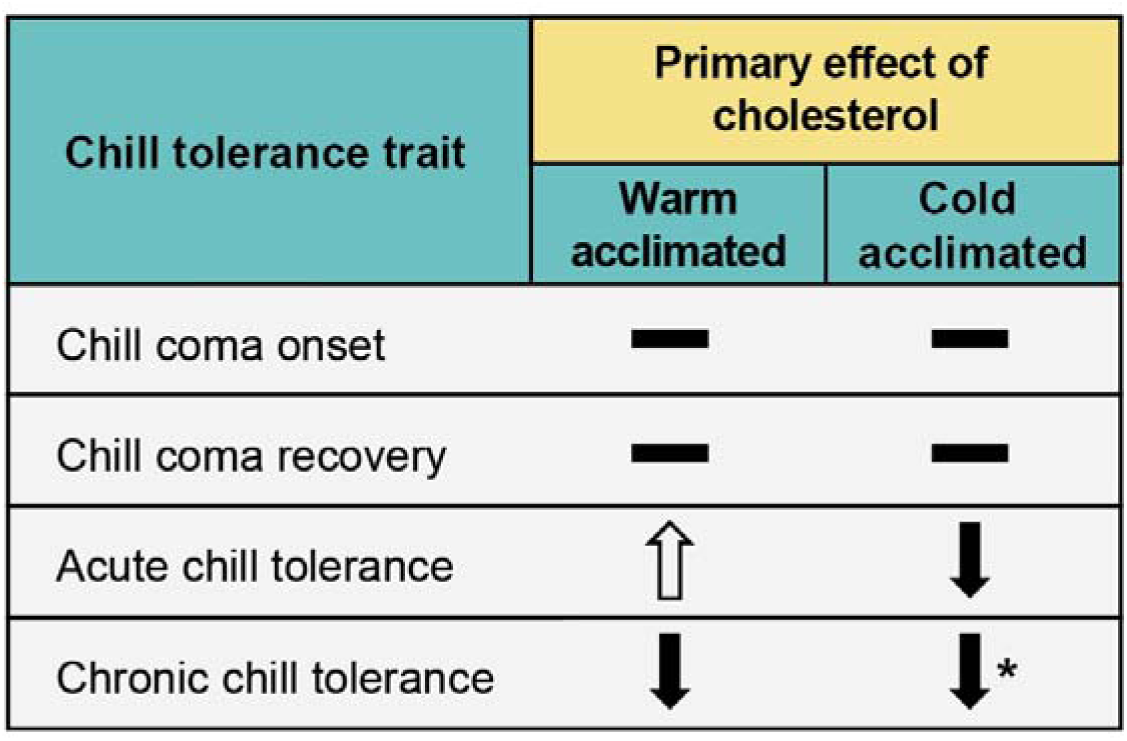
Summary of the main effects of a high cholesterol diet on chill tolerance metrics compared to a control population in *Drosophila melanogaster*.

## Discussion

The effects of a high cholesterol diet on *D. melanogaster* chill tolerance are highly-context dependent. Consistent with previous findings (Shreve et al., 2007), flies raised at relatively warm temperatures (25°C) and fed a high cholesterol diet were more tolerant of an acute cold exposure (Fig. 2). To extend from this observation, we also investigated survival/injury following chronic cold exposures to a milder temperature. Unlike acute chilling, we found the high cholesterol diet had a negative effect on survival following chronic chilling (Fig. 3). The contrast between these results supports the idea that acute and chronic cold exposures cause injury through different mechanisms, and that a high cholesterol diet can help to prevent direct, but not indirect, chilling injury. Generally, it is thought that cholesterol improves cold tolerance by increasing cell membrane fluidity, thereby mitigating phase transitions of membrane phospholipids to the gel phase resulting in mechanical injury to the membrane and subsequent cell death (direct chilling injury) (White, 1993). While acute cold exposure is more often associated with direct injury, it is reasonable to anticipate that high cholesterol feeding would improve tolerance to chronic cold exposure as well. Cholesterol plays a role in modulating the activity of passive and active ion transport, which is known to play an important role in indirect chilling injury. Cholesterol has an inhibitory effect on most types of ion channels, including the majority of potassium channels (Bolotina et al., 1989; Levitan et al., 2014). This effect could help mitigate ion imbalance and extracellular potassium transport at low temperatures or worsen ionoregulatory collapse during chilling. Increased levels of membrane cholesterol increase the activity of Na/K ATPases (Cornelius et al., 2003), which should help maintain ion balance by supporting renal function during chiling (Andersen and Overgaard, 2020; Maddrell and O’donnell, 1992; Yerushalmi et al., 2018). Considering the demonstrated association between chronic chill injury and ion imbalance (Des Marteaux and Sinclair, 2016; Macmillan et al., 2015), our results showing deleterious effects of high cholesterol feeding on chronic cold tolerance were unexpected and emphasize the complexity of mechanisms underlying thermal plasticity.

While data from the survival assays assists in determining the effects of cholesterol on *D. melanogaster* cold tolerance, measuring CCO and CCRT provides additional insight. Employing multiple cold tolerance metrics is useful as variation in one metric is not always associated with variation in others (Davis et al., 2021). We found no effect of a high cholesterol diet on either CCO or CCRT in *D. melanogaster* (Fig. 4). These results suggest that cholesterol has no effect on the ability of flies to maintain neuromuscular function during cold exposure, nor the fly’s ability to recover ion homeostasis after the stress is alleviated. This further supports the idea that cholesterol’s benefit to cold tolerance is exclusive to preventing direct injury to cell membranes as observed in the survival scores.

Unlike cholesterol feeding, but in agreement with prior reports, cold acclimation independently improved all of the measured cold tolerance metrics (Colinet and Hoffmann, 2012; Koštál et al., 2011). While cold acclimation independently improved cold tolerance, it interacted with cholesterol in a way we did not expect. A high cholesterol diet negatively impacted survival of cold acclimated flies following both acute and chronic cold exposures (Fig. 3), suggesting that acclimated flies are more susceptible to both direct and indirect injury when feeding on a high cholesterol diet. We find it difficult to provide a mechanistic explanation for this negative interaction based on the presently available data. One possibility is that additional cholesterol in cell membranes interferes with homeoviscous adaptation during cold acclimation, specifically by interfering with the insertion of phospholipids and fatty acids into the membrane that would otherwise increase membrane fluidity at low temperatures (Michaud and Denlinger, 2006; Overgaard et al., 2008). Analysis of membrane composition following a similar experimental design to our own would allow for a direct test of this hypothesis.

The effects of cholesterol on cold tolerance depend not only on the nature of the cold stress experienced, and prior thermal history, but also on sex. Warm-acclimated female flies had a much steeper decline in survival following both acute and chronic cold exposures (Fig. 2; Fig. 3). Female fruit flies have previously been found to have lower acute cold tolerance based on survival, as well as lower CT_min_ (MacMillan et al., 2009; Parker et al., 2021), although to our knowledge, no efforts have been made to investigate the differences in long term recovery between sexes. We found the opposite effect when examining flies following the chronic cold exposure, where female flies scored higher overall than males, especially when cold acclimated (Fig. 3C, D). Cold acclimation also reduced the sharp decrease in survival in females following both acute and chronic exposures (Fig. 3A, C). Male and female Drosophila therefore appear to substantially differ in their post-chilling survival patterns, specifically in the 24h hours immediately following removal from the cold, and this phenomenon is dependent on acclimation status.

### Conclusions

Feeding on a high cholesterol diet had mixed and unexpected effects on cold tolerance in *D. melanogaster*. The benefits of a high cholesterol diet appear to be exclusive to warm acclimated flies experiencing acute cold exposure, the same conditions under which its positive effects were first reported. Cholesterol supplementation instead had a negative effect on the survival of cold acclimated flies following and acute cold exposure (direct), as well as flies experiencing chronic cold exposure (indirect), regardless of acclimation status. No significant differences were observed in other common cold tolerance metrics (CCO or CCRT). Our findings support the idea that acute and chronic chilling are associated with different mechanisms of injury and highlight the complex manner in which diet can shape thermal tolerance in insects.

## Acknowledgements

The authors would like to thank Hannah Davis for assisting in development of ideas. We would also like to thank Mads Kulmann Andersen for helpful discussion and providing insight throughout the completion of the research.

## CRediT Author Statement

**Mitchell C. Allen:** Conceptualization, Methodology, Validation, Formal Analysis, Investigation, Data Curation, Writing – Original Draft, Writing – Review & Editing, Visualization. **Marshall W. Ritchie:** Conceptualization, Methodology, Validation, Formal Analysis, Investigation, Data Curation, Writing – Review & Editing. **Mahmoud I. El-Saadi:** Conceptualization, Investigation, Methodology, Validation, Formal Analysis, Writing – Review & Editing. **Heath A. MacMillan:** Conceptualization, Methodology, Validation, Resources, Writing – Review & Editing, Visualization, Supervision, Project Administration, Funding acquisition.

## Data availability

All data is provided as a supplementary file for review, and the same file will be included as supplementary material should the manuscript be accepted for publication.

