## Supplemental Figures and Tables for "Effects of a high cholesterol diet on *Drosophila* chill tolerance are highly context-dependent"

**Table S1**: Output from ordinal logistic regression testing the effects of cholesterol supplementation (diet), sex, exposure temperature, and recovery time (and retained interactions) on injury scores following acute cold stress (-4°C or -5°C) in *Drosophila melanogaster*. Analysis was conducted using flies from the 4^th^ generation of high-cholesterol or Bloomington (control) feeding. Odds ratios (OR) and 95% confidence intervals (95% CI) were calculated for factors in brackets. Main effects and interactions in boldface were statistically significant (P < 0.05).

| **Factor** | **OR** | **95% CI** | ***P*** |
| --- | --- | --- | --- |
| Sex­ (male) | 1.98 | 0.68, 5.84 | 0.212 |
| **Diet (cholesterol)** | **4.64** | **2.02, 11.29** | **< 0.001** |
| **Recovery time (24 h)** | **0.01** | **< 0.01, 0.03** | **< 0.001** |
| Exposure temperature (-4˚C) | 1.56 | 0.70, 3.53 | 0.276 |
| Sex (male) : Diet (cholesterol) | **3.29** | 0.80, 15.54 | 0.110 |
| **Sex (male) : Recovery time (24 h)** | **40.79** | **10.35, 169.35** | **< 0.001** |
| Exposure Temperature (-4˚C) : Sex (male) | 3.38 | 0.95, 12.70 | 0.073 |

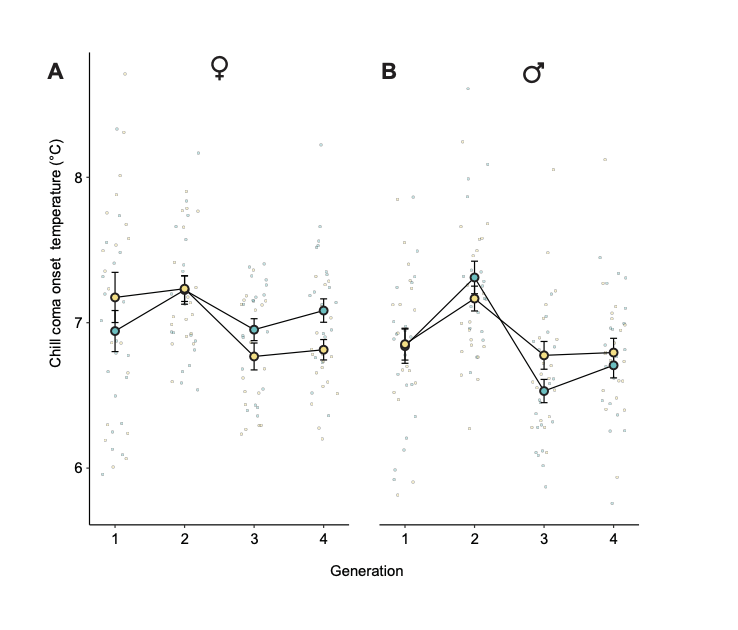

**Figure S1:** **Feeding on a high-cholesterol diet has no clear effect on chill coma onset over four generations.** Chill coma onset temperatures fed on a control or cholesterol-augmented diet. Output from generalized linear model are presented in Table S2. Large points with error bars represent the mean ± sem, and small points represent scores of individual flies. Error bars that are not visible are obscured by the symbols.­­

**Table S2**: Output from generalized linear model testing the effects of generation, cholesterol supplementation (diet), and sex (and retained interactions) on chill coma onset temperature (CCO) in *Drosophila melanogaster*. Main effects and interactions in boldface were statistically significant (P < 0.05).

| **Factor** | **F** | ***df*** | ***P*** |
| --- | --- | --- | --- |
| **Diet** | **< 0.01** | **1,304** | **0.048** |
| Sex | 7.49 | 1,303 | 0.498 |
| Generation | 10.21 | 1,302 | 0.717 |
| Diet : Sex | 1.02 | 1,301 | 0.098 |
| **Diet : Generation** | **1.54** | **1,300** | **0.014** |
| Sex : Generation | 0.25 | 1.299 | 0.053 |
| **Diet : Sex : Generation** | **5.17** | **1,298** | **0.024** |
